# A Cost-effective Purification Process for Erythropoietin Biosimilar through Downstream Process Engineering

**DOI:** 10.1101/2023.01.18.524508

**Authors:** Kakon Nag, Md. Enamul Haq Sarker, Samir Kumar, Sourav Chakraborty, Habiba Khan, Md. Jikrul Islam, Md. Maksusdur Rahman Khan, Md. Mashfiqur Rahman Chowdhury, Rony Roy, Ratan Roy, Bipul Kumar Biswas, Md. Emrul Hasan Bappi, Mohammad Mohiuddin, Naznin Sultana

**Author notes:** Contributed equally (Md. Enamul Haq Sarker, Samir Kumar).

## Abstract

Well-characterized and scalable downstream process for purification of biologics is extremely demanding for delivering quality therapeutics to patients at a reasonable price. Erythropoietin (EPO) is a blockbuster biologic with diverse clinical applications but its application is limited to financially well-off societies due to high price. The high price of EPO is associated with the technical difficulties related to the purification challenge to obtain qualified product with a cost-effective defined process. Though there are reports for purification of EPO but there is no report of well-characterized downstream process with critical process parameters (CPPs) that can deliver EPO consistently satisfying the quality target product profile (QTPP), which is a critical regulatory requirement. To advance the field, we applied quality by design (QbD) principle and design of experiment (DoE) protocol to establish an effective process, which is scalable up to 100× batch size satisfying QTPP. We have successfully transformed the process from static mode to dynamic mode and validated. Insignificant variation (p> 0.05) within and between 1×, 10× and 100× batches showed that the process is reproducible and seamlessly scalable. The biochemical analysis along with the biofunctionality data ensures that the products from different-scale batches were indifferent and comparable to a reference product. Our study thereby established a robust and scalable downstream process of EPO biosimilar satisfying QTPP. The technological scheme presented here can speed-up the production of not only EPO but many other life-saving biologics and make them available to mass population at a reduced cost.

## 1. Introduction

Erythropoietin (EPO) is a glycoprotein cytokine, also known as haematopoietin or haemopoietin [1]. This biomolecule involved in the differentiation, proliferation and maintaining of the physiological levels of erythroid stem cell [2,3]. In adult stage, it is produced by interstitial fibroblasts in the kidney; it is also produced in perisinusoidal cells in the liver at fetal and perinatal period. Recombinant human erythropoietin (rhEPO) is a biotechnologically produced therapeutic protein. EPO and rhEPO are used interchangeably in this article hereafter. EPO is composed of 165 amino acids glycoprotein and its estimated molecular weight is 34 kDa [4]. It has 5 – 8 isomeric forms over the isoelectric point (pI) range of 4.4 – 5.2 [5]. There are 3 potential N-linked glycosylation sites on Asn24, Asn38, Asn83, and 1 O-glycosylation site on Ser126 [6 – 9]. These glycosylations cover approximately 40% molecular weight of the protein. EPO has two disulfide bonds between the cysteines 7 – 161 and 29 – 33, which are essential for maintaining biological activity [4,10]. Functional EPO cooperates with various other growth factors such as interleukin (IL) -3, IL-6, glucocorticoids, and stem cell factor (SCF) to develop erythroid lineage from multipotent progenitors. EPO binds to the EPO receptor (EPOR) on the surface of the RBC progenitor and activates JAK2 signaling cascade [11,12]. This initiates the STAT5, PIK3 and Ras MAPK pathways, which results in differentiation, survival and proliferation of the erythroid cell [13].

EPO has been used to treat anemia related to chronic kidney disease (CKD), cancer chemotherapies, zidovudine patients with HIV-infection etc. [14]. It has been reported that rhEPO is potent to ribavirin treatment in patients with Hepatitis C [15]. CKD is a rising global health problem, and anemia is a serious complication of CKD that has significant adverse outcomes [16]. There are more than 850 million people globally who suffer from CKD, which is increasing every year [17]. More than 37 million American adults may have CKD, and it is estimated that more than 1 out of every 7 people with kidney disease have anemia [18]. It was also reported that study-based estimated prevalence of anemia was 15% among the CKD patients in the USA, 45 – 55% among the Asian CKD patients, and 50 – 90% among the African CKD patients [19].

Rising prevalence of anemia due to CKD has been driving the global demand for the EPO drugs. In 2019, the global EPO drug market was expected to grow more than US $ 18.67 billion by 2025 at a compound annual growth rate (CAGR) of 11.65% during the forecast period (2018-2025) [20]. Raising demand and limited supply of qualified rhEPO lead the treatment of anemia and other blood deficiency diseases expensive. The cost of treatment of anemia by rhEPO was estimated at US $24,128.03 for the Hb level 9 – 10g/dl and US $28,022.33 for the Hb level 11 – 12g/dl per quality-adjusted life-year [21]. In 2012, Matti *et al*., has reported that weight-based dosing cost of biosimilar EPO treatment was €5,484 for once a week, and it increases up to €7,168 [22]. According to a recent report of World Bank, the global average GDP per capita was US $11,570 [23]. Ironically, the average GDP of the global population is below the average medication cost of EPO treatment. Further, the cost of EPO treatment is creeping up steadily due to the growing concerns for regulatory issues associated with the satisfying of the finer-level of the quality target product profile (QTPP). Therefore, it is an ever-growing challenge to ensure quality medication at an affordable price globally.

Bioprocessing of recombinant proteins require a complicated manufacturing process with multiple unit operations. Therefore, it is a highly rewarding task to establish a qualified process that produces cost-effective quality product. Most biotechnology unit operations are complex in nature with numerous process variables and diverse attributes of feed materials, which can have significant impacts on the performance of the process and the QTPP of the final product. The diversity of biotherapeutics and protein expression systems demonstrated the necessity of competent process development that can increase the cost-effective productivity at commercial scale [24]. Quality by design (QbD) using design of experiments (DoE)-based approach offers suitable solutions to this conundrum, and allows for efficient estimation and identification of critical process parameters (CPPs) [25]. The resultant process provides controllable operational conditions for obtaining products satisfying QTPP on regular basis – batch after batch. Therefore, systematic approach (Scheme 1) of process development is a prominent task to achieve QTPP for each biopharmaceutical.

Though many literatures are available regarding protein purification however, to the best of our knowledge, there is none with the definitive process characterization indicating CPPs, and controllability of CPPs to achieve QTPP with the scaleup opportunity. Scalability is one of the vital factors for protein-therapeutic industry. Even though many literatures described the proof of purification for recombinant protein like EPO but they did not provide adequate information and data whether their process is scalable and controllable (Table 1). In addition, validation is an integral part of a process; it involves the systematic study of systems, facilities and processes with the target of determining whether they perform their intended functions properly and consistently as specified. A validated process should be capable of providing a high degree of assurance that uniform batches will be produced and relevant produces shall meet preset specifications. Validation itself cannot improve processes but confirms that the processes have been properly developed and are under control. Moreover, if the process is developed with appropriate CPPs then validation will confirm that the process is under ‘total control’.

**Table 1:** Different purification processes for EPO with their pros and cons.

| References | Process steps | Process performance | Limitations |
| --- | --- | --- | --- |
| Lai and Strickland *et al.* (1985) [28] | AEC → RPC → SEC | Not mentioned. | Incomplete and uncharacterized process, SEC is not economic & suitable for scale up |
| Beck and Withy (1988) [29] | CEC → RPC → SEC | Total yield (%): 60 → 50 → 41 | Incomplete and uncharacterized process. Need to adjust sample pH below 4 resulting precipitation or degradation, SEC is not economic & suitable for scale up |
| Sugaya *et al.* (1997) [30] | AFC → AFC → SEC | Total yield (%): 79 → 54 → 40 | Incomplete and uncharacterized process, SEC is not economic & suitable for scale up |
| Zanette *et al.* (2003) [31] | AFC → AEC → SEC | Total yield (%): 76 → 60 → 30 | Incomplete and uncharacterized process. Need to adjust sample pH below 4, resulting precipitation or degradation, SEC is not economic & suitable for scale up, Impurities were not removed based on hydrophobicity |
| Hu *et al.* (2004) [32] | AEC → HIC → SEC | Total yield (%): 51 → 46 → 43 | AFC is more efficient to enrich glycosylated protein compare to AEC, SEC is not economic & suitable for scale up, AEC is more suitable to separate charge variant compare to HIC |
| Carcagno *et al.* (2006) [33] | HIC → AEC → CEC → SEC | Total yield (%): 70 → 56 → 40 → 30 | Need to add high molar salt at first step, SEC is not economic & suitable for scale up, need extra process step to complete the process |
| Goletz and Stöckl (2012) [34] | RPC → AFC → SEC → AEC | Final host cell protein content less than 0.01% | AFC is more efficient to enrich glycosylated protein compare to RPC, SEC is not economic & suitable for scale up and not suitable after AFC, |
| Wojchowski *et al.* (1987) [35] | HIC → AFC | Total yield (%): 100 → 55 | Incomplete and uncharacterized process, not suitability in production scale |
| Ghanem *et al.* (1994) [36] | AFC → AEC | Total yield (%): 73 → 53 | Incomplete and uncharacterized process, not suitability in production scale |
| Hsu and Chang (2002) [37] | AFC → SEC | in vitro activity: 240 kU/mg | Incomplete and uncharacterized process, not suitable for scale-up, Yield was not claimed |
| Surabattula *et al.* (2011) [38] | AFC → AEC | Final total yield: 42% | Incomplete and uncharacterized process, not suitable for scale-up |
| Broudy *et al.* (1988) [39] | AFC → AEC → RPC | Total yield (%): 65 → 49 → 35 | After RPC, acetonitrile removal is difficult to formulate the sample, need other steps which are not performed and characterized |
| Quelle *et al.* (1989) [40] | AEC → RPC → AFC | Total yield (%): 95 → 85 → 80 | AFC is more efficient to capture glycosylated protein, difficult to formulate the sample after AFC, need other steps which are not performed and characterized |
| Merrifield (1990) [41] | AFC → AFC → AEC | Total yield: 80% (in the first step) | Need to add extra TFF process before each chromatography step, difficult to formulate the sample after AFC, need other steps which are not performed and characterized |
| Burg *et al.* (2002) [42] | AFC → HIC → HAC → RPC → AEC | Final total yield: 25% | Need other steps to formulate the sample which are not performed and characterized, After RPC, elute is not suitable to proceed for AEC. |
| Koh *et al.* (2014) [43] | AEC → HAC → AEC | Total yield (%): 60.1 → 54.2 → 18.1 | AFC is more suitable for enrichment of glycosylated protein, need other steps which are not performed and characterized |
| Bandi *et al.* (2017) [44] | AEC → MMC → AEC → CEC → AFC → AEC → SEC | Purity (%): ≥ 98% | Need at least three TFF process steps and other steps to formulate the sample which are not characterized, too many process steps increase process cost, SEC is not economic & suitable for scale up, Specific yield was not claimed |

Biopharmaceutical industries can meet the QTPP and regulatory requirements by achieving a validated process embedded with well-characterized and defined CPPs [26, 27]. Henceforth, it is extremely important to establish such a process for cost-effective EPO manufacturing which is scalable, controllable and validated for satisfying QTPP, and will ultimately meeting regulatory requirements. Here, we report such a robust and scalable EPO purification process embedded with defined CPPs and meeting QTPP at a lower risk and reduced cost.

## 2. Materials and methods

### 2.1 Screening of binding and elution conditions of EPO

Different chromatography resins i.e., Capto blue affinity chromatography resin, Q Sepharose anion exchange chromatography resin, Source reversed phase chromatography resin, and MacroCap SP cation exchange resin (GE Healthcare, USA) were selected, and different buffer conditions were applied as mentioned elsewhere in the manuscript to identify suitable condition for each resin to capture and elution of target product with reduced impurities. Approximately 100 μl of each resin were taken in 96-well plate during screening. Each resin was activated with equilibration buffer, and 1ml of filtered (0.6 μm followed by 0.2 μm PES) sample was applied for interaction with resin. Samples were washed with equilibration buffer followed by elution with elution buffer. Buffer of AFC eluate was then exchanged by AEX equilibration buffer using 10 kDa NMWCO Vivaspin (Sartorius Stedim, Germany) column then applied in AEX resin; after wash with equilibration buffer, the products were eluted with elution buffer. AEX eluate was then applied to RPC resin, and the desired product was eluted after wash with equilibration buffer. RPC eluate was then applied to CEX resin; after washing of the resin with equilibration buffer the products were eluted in elution buffer. The flow through, wash, elution and cleaning in place (CIP) samples of each chromatography resin were tested by dot blotting to identify the desired product. CIP was performed with 1N NaOH solution for the AFC and AEX resins, whereas 0.5N NaOH solution was used for RPC and CEX resins.

### 2.2 DoE for the unit processes of purification train

DesignExpert 13 software (Stat-Ease Inc., USA) was used for DoE. CPPs and optimum buffer conditions for AFC, AEX, RPC and CEX chromatography were identified using 2-level factorial optimal (Custom) design. Data were analyzed using surface response plot. Combinations of pH levels and NaCl concentrations for AFC (Supplementary Taable 1) and AEX (Supplementary Table 2), pH and aquous:organic solvent for RPC (Supplementary Table 3), and pH and CV for CEX (Supplementary Table 4) were analyzed, respectively. The recommended chromatography conditions for each chromatography step were completed in static mode in 1 ml size in microcentrifuge tube. Recovery % of eluted sample of each step for all run conditions were analyzed by analytical size exclusion chromatography (SEC). Recovery data of target product satisfying QTPP for all chromatography steps were processed in DesignExpert 13 software. Optimum operating conditions and CPPs were identified from the surface response plot. All data points were validated using series of triplicate experiments.

### 2.3 Scale-up and adaptation of the unit purification processes in dynamic mode

After finding the binding and elution conditions (design space) of product, the unit purification process steps were scaled up for the adaptation in dynamic mode from static mode. The purification process of EPO was optimized at 1× batch size (50 ml), which was compatible with small scale column formats. The chromatography conditions were adapted through two consecutive small scale (1×) batches and then validated with next three consecutive batches. After adaptation of process in small scale, reproducibility of overall yield percentages among the batches were tested following first degree of statistical approach and the acceptance criteria of those process parameters were set. The sample was sequentially filtered through 0.6 μm and 0.2 μm PES filter (Sartorius Stedim, Germany), and EPO was captured onto equilibrated AFC column (GE Healthcare, USA) in AKTA pure 25 system (GE Healthcare, USA) followed by washing and elution. FPLC systems and columns were sanitized before and after each purification step. FPLC results were evaluated through Unicorn 7.0 software (GE Healthcare, USA) for all chromatography steps. The AFC-eluates were buffer exchanged with 10kDa Vivaspin protein concentrator (Sartorius Stedim, Germany) using Sorval Lynx XTR centrifuge (Thermo Scientific, USA). The protein concentrator was sanitized before and after diafiltration. The retentate was loaded onto AEX column in AKTA pure 25 system. The AEX-eluate was then loaded onto RPC column in AKTA pure 25 system. The column was equilibrated and washed with acidic solution followed by elution in mixed gradient between wash buffer and organic solvent. The eluates from RPC were incubated for virus inactivation at low pH. The virus-inactivated samples were loaded onto CEX column in AKTA pure 25 system. The in-line pH, conductivity as well as sample in-line pH, conductivity and quantity of buffers were considered as in-process check (IPC) points in AFC, AEX, RPC and CEX steps. Residence times for chromatography were maintained at 6 – 10 minutes, and delta column pressure was maintained at below 6 bar. The samples were then passed through 0.22 μm filter followed by 20 nm ViroSart virus filter (Sartorius Stedim, Germany). The clarified samples were then reconstituted in formulation buffer. After passing through 0.45|0.2-micron Minisart sterilizing filter (Sartorius Stedim, Germany), samples were stored at 2 – 8 °C and analyzed.

### 2.4 Scale-up and validation of 10× (500 ml) and 100× (5000 ml) batches

After adaptation of 1× batch in dynamic mode and optimization of operating conditions, the process was scaled up at 10× (500 ml) batch size by extrapolating CPPs for relevant unit operations. The sample was processed for AFC unit process and then subjected onto AEX unit process after completion of tangential flow filtration (TFF). Sartocon Slice TFF cassette (Sartorius Stedim, Germany) and AKTA Flux 6 (GE Healthcare, USA) were used for buffer exchange in this unit process step. AEX eluates were subjected for RPC unit process followed by virus inactivation. Samples were then processed for CEX unit process and reconstituted in formulation buffer. All IPC parameters were applied in 10× batches. Three consecutive 10× batches were performed to validate the process, and samples were analyzed for QTPP. Similarly, all unit processes were further scaled up to 100× batch size (5000 ml) maintaining IPCs and CPPs, and samples were analyzed for QTPP. Appropriate quantity of resins, relevant columns, hardware and buffers were used for successive scale-up of unit processes.

### 2.5 Analytical approach for QTPP confirmation

#### 2.5.1 Dot blotting

PVDF transfer membrane (0.2 μm; Thermo Fisher Scientific, USA) was cut based on sample number and regenerated in methanol. The membrane was equilibrated in transfer buffer (pH 8.3). Subsequently, 10 μL of each sample was loaded on the membrane, and allowed to dry. The reactivated membrane was blocked and treated anti-Epo polyclonal antibody (Thermo Fisher Scientific, USA). Goat anti-rabbit (H+L) IgG HRP conjugated (Thermo Fisher Scientific, USA) was used as secondary antibody. Novex^®^ ECL Chemiluminescent Substrate (Thermo Fisher Scientific, USA) for HRP was used to detect the signal, and the images were captured using Amersham Imager 600 RGB (GE Healthcare, USA).

#### 2.5.2 Particle size distribution

Samples were prepared in 0.22-micron filtered 1× PBS (pH 7.2), and after stabilization at 20 ºC for 20 min, analyzed in disposable plastic cuvette using a Zetasizer Nano ZSP (Malvern Panalytical Ltd., UK) where respective buffers used as dispersant. The equipment was switched on minimum 30 – 60 minutes before the experiment to stabilize the system. The refractive index (RI), viscosity and dielectric constant of dispersion buffer (1× PBS, pH 7.2 at 20 °C) was considered 1.33, 0.88 cPs and 79, respectively.

#### 2.5.3 Western blotting

The Western blots were performed following standard operating methods. In short, samples were processed individually in microfuge tubes. Laemmli buffer was used at 5× dilution, denatured by heating for 3 minutes at 95ºC, cooled on ice, and 25 μl of each sample was loaded into a commercial gel (Thermo Fisher Scientific, USA). After the gel electrophoresis, samples were transferred onto activated PVDF membrane (Thermo Fisher Scientific, USA). The EPO was detected and visualized using specific antibodies (Thermo Fisher Scientific, USA) and chemiluminescence. The signals were captured using Amersham Imager 600 RGB (GE Healthcare, USA).

#### 2.5.4 Chromatography for determination of assay and impurities

The assay for EPO of the samples were determined using Vanquish UHPLC system (Thermo Fisher Scientific, USA). In short, the samples and reference product Eprex^®^ (20 μL of each) were applied in RPC column (Hypersil GOLD C8, 175 Å, 2.1× 100 mm, 1.9 μm column) (Thermo Fisher Scientific, USA) for analysis. The formulation buffer was considered as base line reference. A gradient of mobile phase A (water with 0.1% FA) and mobile phase B (90% ACN in water with 0.1% FA) was used as carrier solvent at a flowrate of 0.3 mL/min. The impurity profiles of the samples were analyzed using SEC in reference with Eprex^®^. Analyses were performed in an Ultimate 3000 RSLC system (Thermo Fisher Scientific, USA) using 50 μL samples in a Biobasic SEC-300 column (300 mm*7.8mm, 5μm) (Thermo Fisher Scientific, USA). Phosphate buffer (pH 7.4) was used as mobile phase at a flow rate of 1.0 ml/min. The column temperature was maintained at 70 °C, and run time was 25 minutes for both methods. The signals were detected at 280 nm, and chromatograms were recorded.

#### 2.5.5 In vitro functional assay using cell culture

The *in vitro* functional assay was performed using the TF-1 cell line, which was originally derived from a patient diagnosed with erythroleukemia. TF-1 cells were maintained in RPMI 1640 medium (Gibco, USA) supplemented with 10 % FBS (Thermo Fisher Scientific, USA), 1% PS (Thermo Fisher Scientific, USA), and 5 ng/mL human recombinant GM-CSF (Thermo Fisher Scientific, USA) at 37 °C and 5 % carbon dioxide. Cells were washed twice in PBS to eliminate GM-CSF, and then were seeded at a density of 10^5^ cells/well in a 24-well tissue culture non-treated cell-culture plate. Cells were grown for 72 h in the presence or absence of originator (Eprex®) and EPO samples from each batch at indicated concentration. Cells were collected, centrifuged, re-suspended in 1 mL of PBS, and counted by Countess 2 automated cell counter (Thermo Fisher Scientific, USA).

#### 2.5.6 Sterility and endotoxin testing

Bacterial and fungal sterility were tested using direct inoculation technique. Samples (1 mL) were inoculated in 10 mL of Tryptic Soya Broth (Sigma Aldrich, USA) media for 14 days and absorbance were measured at 600 nm. Endotoxin in samples were tested using Pierce LAL Chromogenic Endotoxin Quantitation Kit (Thermo Fisher Scientific, USA) as per supplier’s instructions.

## 3. Results

The study design and management process were performed according to the decision tree (Scheme-1A) and process flow diagram (Scheme-1B). The CPPs were identified according to the schema. Iterative applications of the relevant steps were followed when it was necessary.

### 3.1 Screening of binding and elution conditions of EPO

After screening of AFC resins, we found that 20 mM Tris-HCl, pH 7.4 was suitable for equilibration and washing for the selected matrix, and 1.5 M NaCl in wash buffer was appropriate for elution of the sample. After dialysis against Tris buffer at neutral pH, the conductance and pH were found ≥ 3.0 mS/cm and 7.0, respectively. For AEX chromatography, we observed that similar buffer composition of AFC at neutral pH was suitable for binding and elution of EPO. In the RPC chromatography process, 0.1% of TFA in WFI was found appropriate for resin equilibration and wash, and the product was eluted in 95% of acetonitrile. In CEX chromatography process, 20 mM Glycine, pH 2.0 was found suitable for resin equilibration and wash. Sodium phosphate buffer with 150 mM NaCl, pH 7.20 was effective for eluting the EPO satisfying QTPP. Representative samples of different process steps were identified with dot blot analysis. (Figure 1).

**Figure 1:**
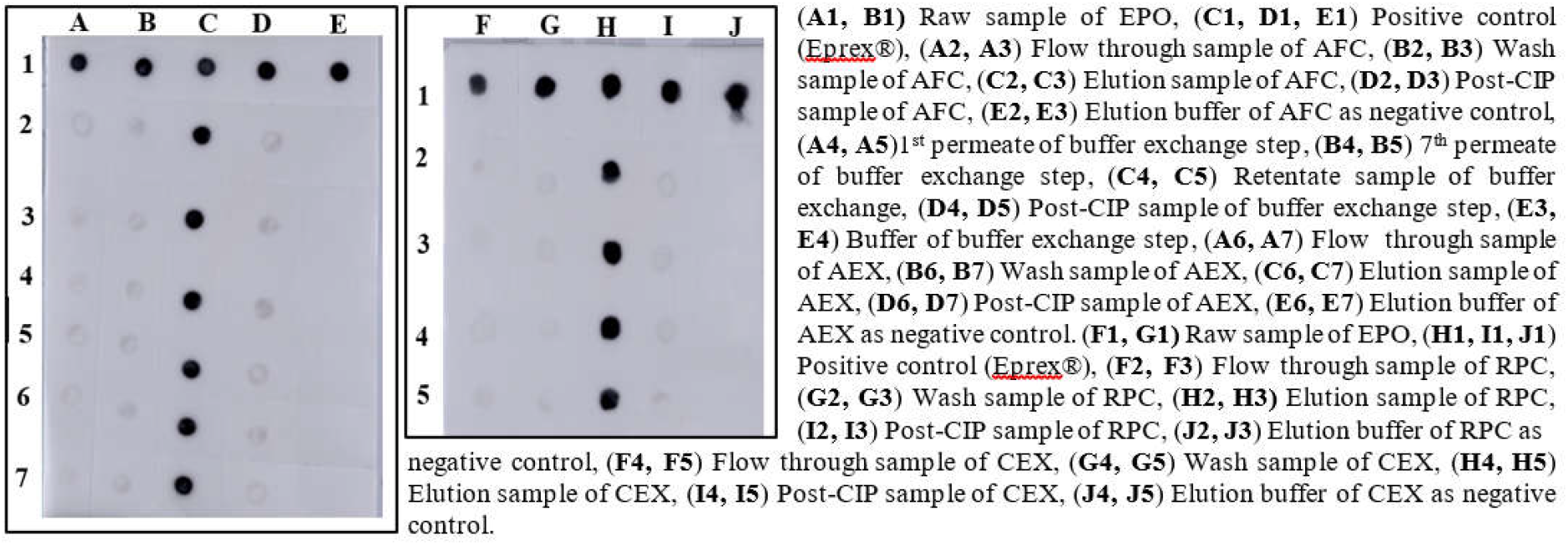
Screening of binding and elution conditions of EPO using different resins.

### 3.2 Design of experiments (DoE) of purification process

From contour plot and 3D surface response plot, the pH range of media for the AFC unit process was extrapolated within 6.7 – 8.4 (control space), more preferably at 7.4 ± 0.2 (operation space). Sodium chloride concentration range was extrapolated within 1400 – 1700 mM (control space), more preferably at 1500 ± 100 mM (operation space), which are equivalent to 120.4 – 146.2 mS/cm and 129 ± 10 mS/cm, respectively. The actual value and predicted value of recovery percentage were within close distribution to the regression line indicates that the pH and NaCl concentration both have strong effect on AFC recovery (Figure 2). The model terms are significant where F-value was found 9.49 and *p* value was 0.0015, and suggested that the model was well-fitting. After analyzing eluted samples of all run conditions, we found less recovery with higher impurities at a pH lower than 6.7 with 1500 mM [NaCl] carrier. In contrast, we have observed adequate recovery with higher impurities at a pH level above 8.4 with 1700 mM [NaCl] carrier (Supplementary figure 1A). The recovery range of target product was found ≥ 80% with the least impurities in control space, and was considered as acceptable limit.

**Figure 2:**
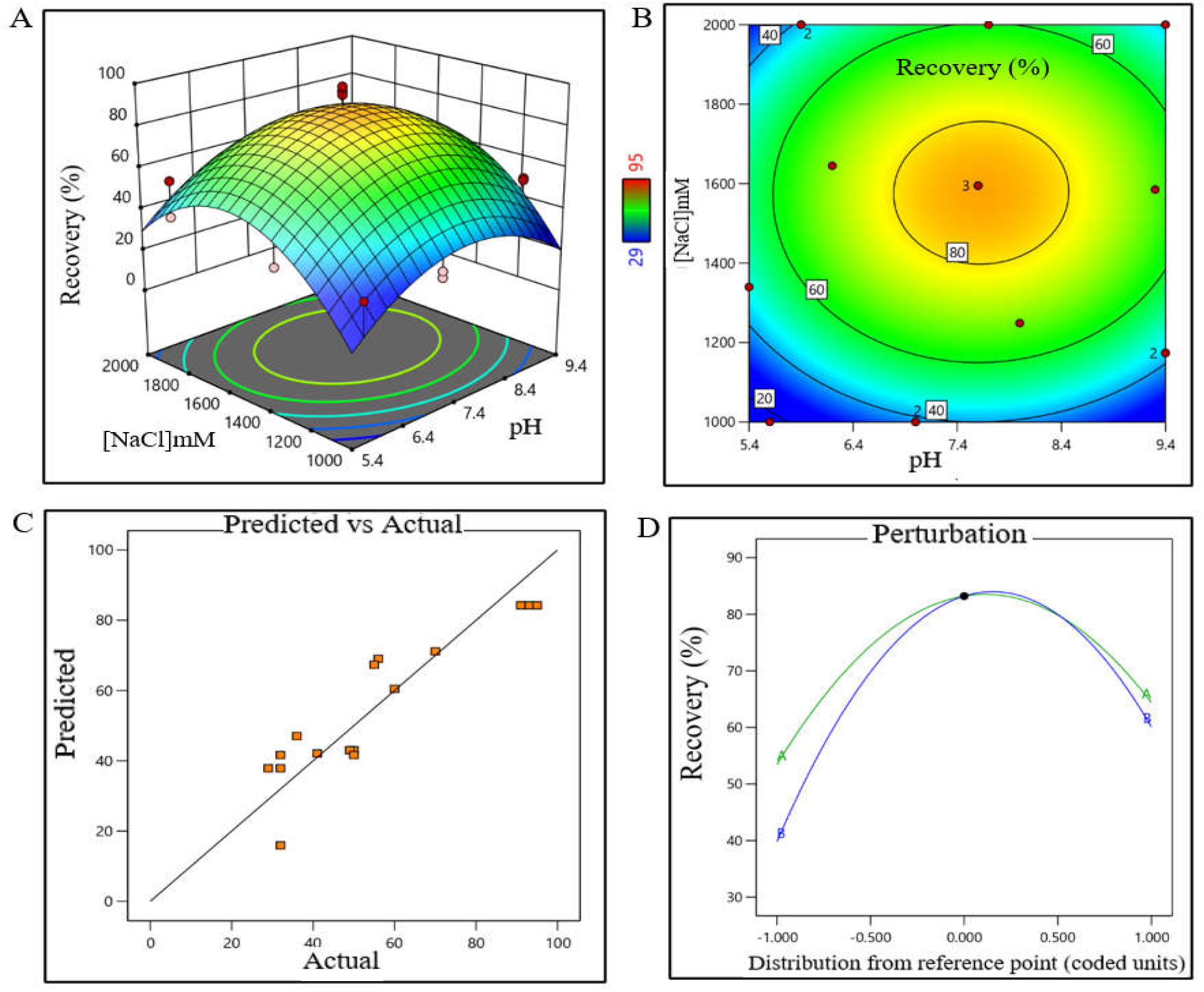
DoE surface response plot of AFC process step in static mode. (**A)** 3D surface response plot of elution condition, (**B**) Contour plot of elution condition, (**C)** Actual *vs* predicted response of recovery %, and (**D**) Perturbation of recovery % against two factors pH and [NaCl].

For the AEX unit process, the buffer pH range was extrapolated within 6.5 – 7.7 (control space), more preferably at 7.0 ± 0.2 (operation space). NaCl concentration range was extrapolated within 260 – 380 mM (control space), more preferably at 300 ± 15 mM (operation space), which are equivalent to 22.5 – 32.70 mS/cm and 25.5 ± 1 mS/cm, respectively. The actual value and predicted value of recovery percentage were in a close distribution to the regression line indicates that the pH and NaCl concentration both have strong effects on AEX recovery within the operation range (Figure 3). The model terms were significant where F-value was found 20.58 and *p* value was <0.0001, and suggested that the model was well-fitting. After analyzing eluted samples of all run conditions, we found less recovery with higher impurities at a pH lower than 6.5 with 260 mM [NaCl] carrier. In contrast, adequate recovery with higher impurities were observed at a pH above 7.7 with 380 mM [NaCl] carrier (Supplementary figure 1B). The recovery range of target product was found ≥ 40% with the least impurities in control space and was considered as acceptable limit.

**Figure 3:**
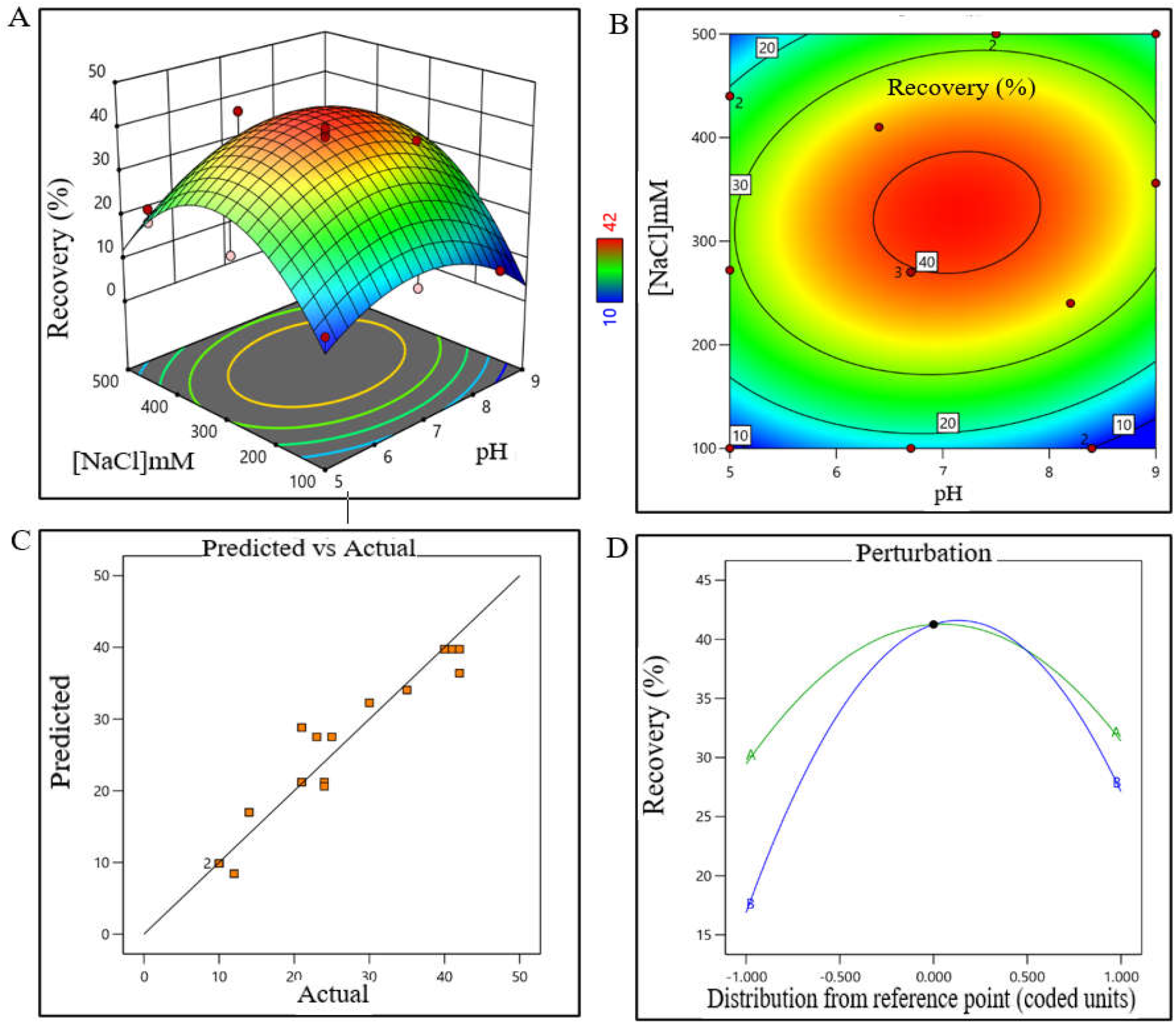
DoE surface response plot of AEX process step in static mode. (**A)** 3D surface response plot of elution condition, (**B**) Contour plot of elution condition, (**C)** Actual *vs* predicted response of recovery %, and (**D**) Perturbation of recovery % against two factors pH and [NaCl].

For the RPC unit process, the pH range was extrapolated within 2.2 – 2.7 (control space), and more preferably at 2.4 ± 0.2 (operation space); whereas acetonitrile concentration range was extrapolated within 45 – 67% (control space), and more preferably at 52 ± 5% (operation space). The actual value and predicted value of recovery percentage were found closer to the regression line within the operation range and suggested that the pH and concentration (%v/v) of acetonitrile both have strong effect on RPC recovery (Figure 4). The model terms were significant where F-value was found 6.20 and *p* value was 0.0072, which have suggested that the model was well-fitting. After analyzing eluted samples of all run conditions, we found less recovery with higher impurities at a condition pH range lower than 2.2 and 45% acetonitrile. In contrast, adequate recovery with higher impurities were observed at a pH range above 2.7 with 67% acetonitrile (Supplementary figure 1C). The recovery range of target product was found ≥ 40% with the least impurity within the control space, and was considered as acceptable limit.

**Figure 4:**
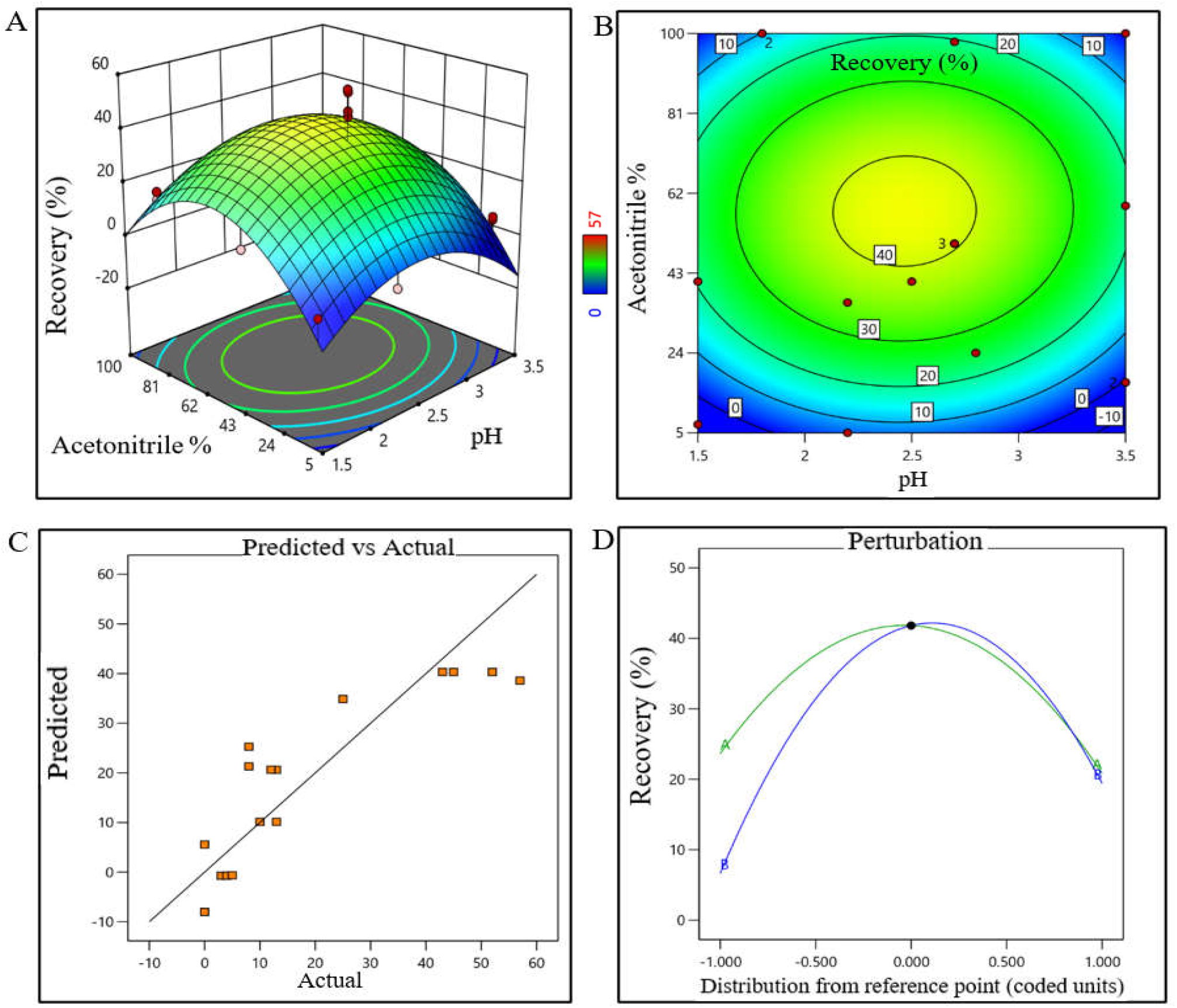
DoE response plot of RPC process in static mode. (**A)** 3D surface response plot of elution condition, (**B**) Contour plot of elution condition, (**C)** Actual *vs* predicted response of recovery %, and (**D**) Perturbation of recovery % against two factors pH and acetonitrile%

The pH range of buffer for CEX unit operation was extrapolated within 4.8 – 6.8 (control space), and more preferably at 5.8 ± 0.2 (operation space) though the plot showing bimodal effect. The buffer volume was extrapolated within 2.5 – 4 CV (control space), more preferably at 3.5 ± 0.2 CV (operation space) to obtain the best outcome. The actual value and predicted value of recovery percentage were found closer to regression line and indicate that the pH and buffer volume (CV) both have strong effects in CEX recovery (Figure 5). The model terms were significant where F-value was found 176.36 with a *p* value <0.0001, which have suggested that the model was well-fitting. After analyzing eluted samples of all run conditions, we found less recovery with impurities at a pH lower than 4.8 with 2.5 CV of buffer volume. In contrast, adequate recovery with higher impurities were observed at a pH above 6.8 with 4 CV of buffer volume (Supplementary figure 1D). The recovery range of target product was found ≥ 80% with the least impurities within the control space, and was considered as acceptable limit.

**Figure 5:**
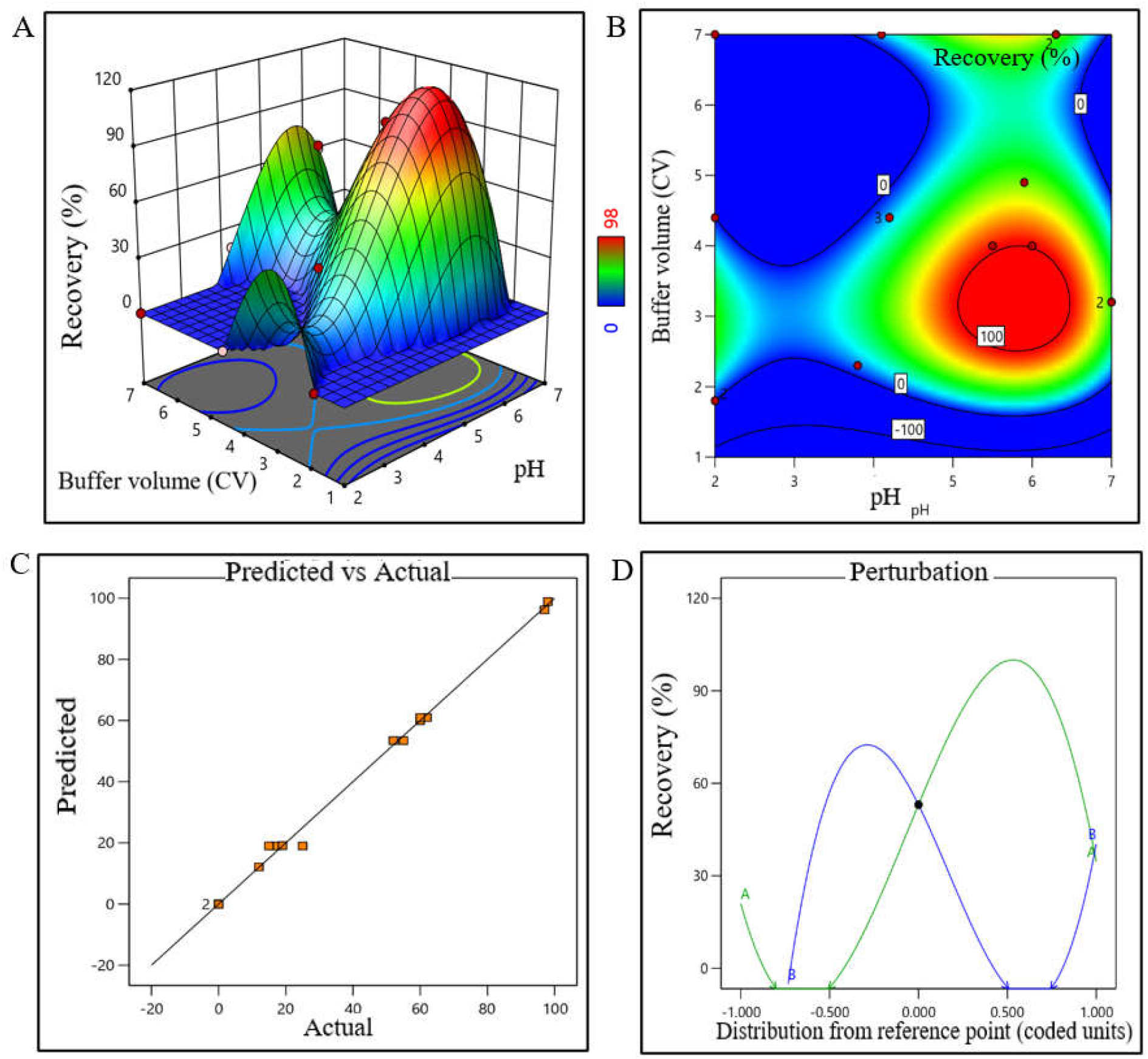
DoE response plot of CEX process in static mode. (**A)** 3D surface response plot of elution condition, (**B**) Contour plot of elution condition, (**C)** Actual *vs* predicted response of recovery %, and (**D**) Perturbation of recovery % against two factors pH and buffer volume (CV).

### 3.3 Adaptation of unit process in dynamic mode, and scale-up

Five consecutive batches were conducted to adapt the process in dynamic mode using chromatography column format and machine-controlled flow conditions, and the acceptance criteria were justified for each unit process steps those were identified from DoE runs. The quantity of the starting materials of these five batches were 45 – 55 mg, the volume were approximately 50 ml and concentration of EPO were 0.9 – 1.1 mg/ml (Supplementary table 5). In case of AFC unit process, it was observed that the pH range of elution buffer and eluted sample were 7.43±0.12 and 7.35±0.14, respectively. The conductivity range (mS/cm) of elution buffer and sample were 123.42±1.61 and 123.78±1.22, respectively. The average eluted protein quantity and yield % were found 41.62±2.38 mg and 81.11±2.66 mg, respectively. The pH and conductance were considered as CPPs and % of yield was considered one of the CQAs for this step (Supplementary table 6). The pH range of retentate sample, conductance (mS/cm), quantity (mg) and yield % were 7.35±0.15, 2.53±0.29, 41.09±2.27 and 98.73±0.31, respectively; where conductance of the retentate sample was considered as CPP (Supplementary table 7). Since, the retentate sample pH, conductivity range and yield percent were not evaluated in DoE (in static mode), the average value and standard deviation of the parameters of these five batches were considered as acceptance limits. In AEX unit process, elution buffer and eluted sample pH were found 7.05±0.14 and 6.97±0.11, respectively; whereas, the conductivity (mS/cm) range were found 123.11±1.33 and 24.09±0.69, respectively. The average quantity of eluted protein was 16.86±1.24 mg with the yield % of 41.01±1.03. The pH of the buffer and the conductance of the eluted sample were considered as CPP; whereas % yield was considered as one of the CQAs for this unit process (Supplementary table 8). The pH, conductivity and yield percent were within the acceptable limit which were identified through DoE. In the RPC unit process, the pH range of elution buffer and eluted sample were observed 2.43±0.09 and 2.17±0.11, respectively. The acetonitrile % of eluted sample was found 50.80±1.92. The average eluted protein quantity and yield % were 8.68±0.57 mg and 51.51±1.72, respectively. The pH of the buffer and % (v/v) of acetonitrile were considered CPPs, and yield % were considered as one of the CQAs (Supplementary table 9). The pH of the elution buffer, % (v/v) of acetonitrile and yield % were within acceptable limit, which were identified as DoE output. After completion of RPC eluted sample was kept 120 minutes to inactivate virus particle and then proceed CEX. For the CEX unit process, it was found that the pH range of elution buffer and eluted sample were 7.07±0.20 and 5.81±0.11, respectively. A 3.5 CV of elution buffer was found sufficient to elute desired product where average eluted protein quantity and yield % were 7.47±1.02 mg and 85.86±7.42, respectively. The pH of the elution buffer and buffer volume were considered as CPPs, and % of yield was considered as one of the CQAs for this step (Supplementary table 10). The pH and the buffer volume (CV) of the elution buffer and yield % were within acceptable limit, which were identified as DoE output. In the virus filtration (VF) step, the pH, conductance (mS/cm), quantity (mg) and yield % were 7.12±0.12, 12.54±0.43, 7.38±0.97 and 98.80±0.65, respectively. The pH of the filtrate sample and time were considered as CPPs, whereas the % of yield was considered one of the CQAs (Supplementary table 11). Since, the % yield was not evaluated in DoE (in static mode), the average value and standard deviation of the parameters of these five batches were considered as acceptance limits. After completion of the sterile filtration (SF) step, it was observed that the pH and conductance of the sample and yield % were similar to the previous step (Supplementary table 12). The overall yield % for the whole process was calculated to 14.14±0.87 % (Supplementary table 13). Since, the overall % yield was not evaluated in DoE (static mode), therefore the average value and standard deviation of the yield % of these five batches were considered as acceptance limits. For all the dynamic unit processes, the CPPs were controlled and monitored using inline automated system.

### 3.4 Validation of unit processes for 10× (500 ml) batch

After completion of purification process optimization, batch size was scaled up to 10× (500 ml) and three consecutive batches (Batch No. 06, 07 and 08) were conducted to validate the purification process. The overall yield percentage of batch no. 06, 07 and 08 were found 13.97%, 14.07% and 13.90 %, respectively, which were within the acceptance range (14.14±1.0%) of yield percentage (Supplementary table 14). The recovery percentage data for individual unit process steps were also within the limit for these three batches (Supplementary table 14). IPC parameters of chromatography steps were monitored and controlled by inline FPLC system during the batch runs.

### 3.5 Validation of 100× (5000 ml)

Batch size was further scaled up to 100× (5000 ml), and three consecutive batches (Batch No. 09, 10 and 11) were conducted to validate the scaled-up process. The overall yield percentage of batch no. 09, 10 and 11 were found 13.30, 14.62 and 13.31 %, respectively, which were within the acceptance range (14.14±1.0%) of yield (Supplementary table 15). The recovery percentage data for individual unit process steps were also within the limit for these 3 batches (Supplementary table 15). All IPC parameter of chromatography steps were monitored by automated inline FPLC system for these batches. The representative chromatogram is shown in Figure 6 (A– D).

**Figure 6:**
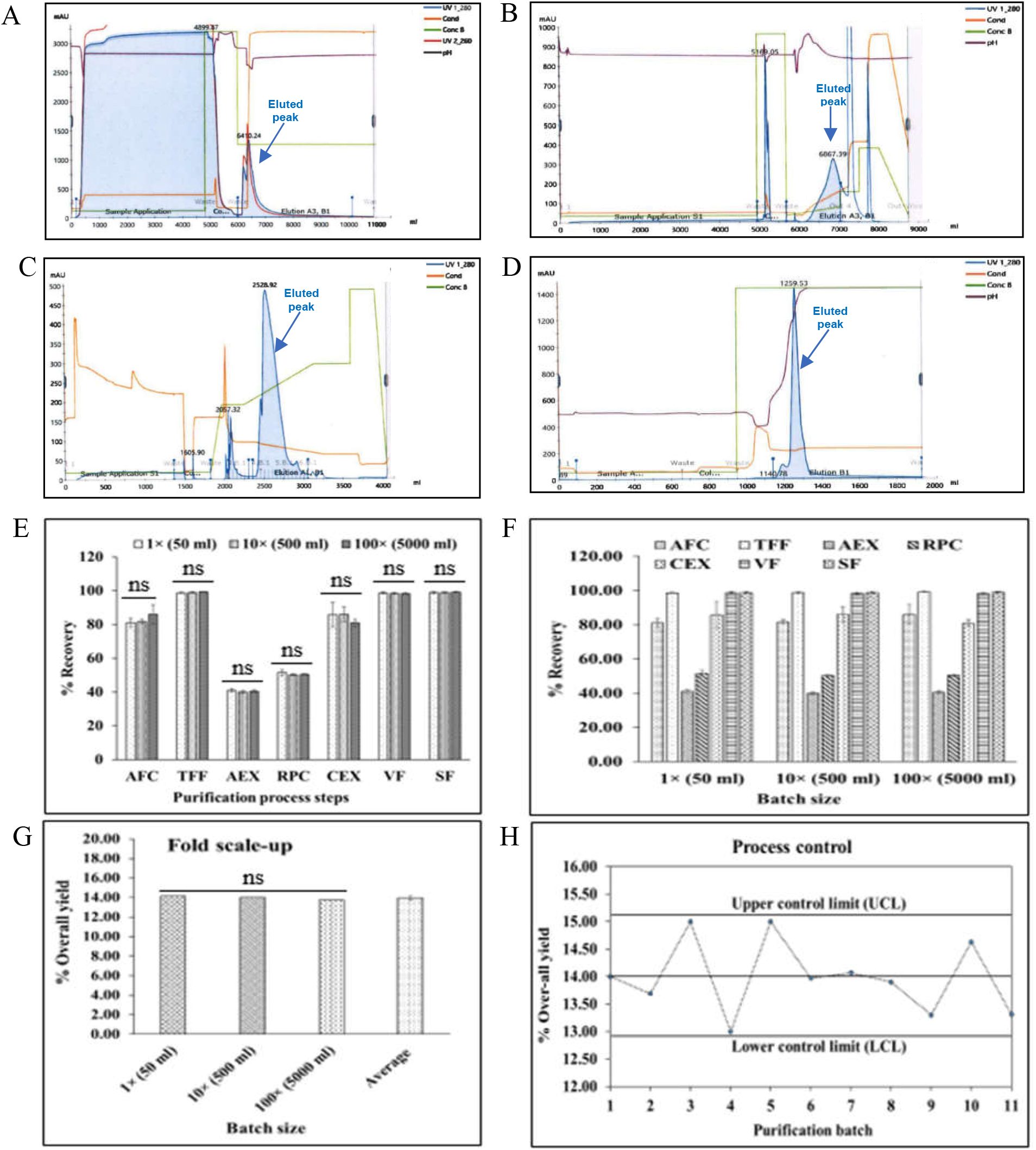
Process monitoring and qualification. (**A**) Representative AFC chromatogram of 100× batch size (**B**) Representative AEX chromatogram of 100× batch size, (**C**) Representative RPC chromatogram of 100× batch size, and (**D**) Representative CEX chromatogram of 100× batch size. **(E)** Recovery % of each step in dynamic mode, and reproducibility between process steps of 1×, 10× and 100× batch size, **(F)** Recovery % of each process step and reproducibility within batches of 1×, 10× and 100× batch size, **(G)** Reproducibility of yield between 1×, 10× and 100× batch, **(H)** Controllability of purification process for 1× (batch 1 – 5), 10× (batch 6 – 8) and 100× (batch 9 – 11) batch sizes, shown in ‘x’ axis.

There were no significant differences observed for the qualitative and quantitative parameters of each unit processes of 1×, 10× and 100× validation batches (Figure 6E, *p*>0.05). We also found that the variation between 3 batches for each batch size (1×, 10× and 100×) is insignificant (Figure 6F, *p*>0.05). The yield percentage of 3 scaled-up batches for both scales (10× and 100×) have no significant variation either (Figure 6G, *p* >0.05). The trend analysis for all these batch’s data for yield % were within the control limit (acceptance range 13 – 15%) (Figure 6H).

### 3.6 Analysis for QTPP

After confirmation of the validation of 1× (50 ml), 10× (500 ml) and 100× (500 ml) batch size, relevant samples were analyzed as per specification, and found all parameters of specification were in accordance within the relevant acceptance limits (Supplementary table 16). The particle size distribution was compared between reference product Eprex® and different batches of GBPD002, where representative results were shown in Figure 7. The representative result for identification test, impurities profile and biofunctionality analysis between reference product Eprex® and GBPD002 were shown in Figure 8. The Western blot data showed that all batches were having same protein species with similar migration pattern (Figure 8A); no secondary bands were observed in the blot. RP-UHPLC analyses data provided clear understanding that the similar quantity of EPO was harvested and were present in each batch (Figure 8B). The SEC-UHPLC result did not show any significant differences between the GBPD002 harvested from different batches and with reference product, suggesting that there is no unwanted species present in any of the EPO preparations (Figure 8C). Comparing the buffer chromatograms, the secondary peaks for both RPC and SEC can be attributed to the buffer and carrier system. All these analyses collectively confirmed the absence of any macromolecular entity that can be generated from process chemistry or product aggregates. The TF-1 cell proliferation data clearly revealed that the materials from different batches and reference responded similarly. Compared with the mock-controlled dishes, which has reduced to half by number from its original seeding population (10^5^/well), all samples assisted the growth of the TF-1 cell population to 3 times of the originally seeded cell numbers (Figure 8D). This data has clearly established that the EPO preparations are similarly active like the reference product. Very close responses for the biofunctionality among the EPO preparations from 3 different batch sizes (1×, 10× and 100×) suggested indifferences among these scaled-up EPO preparations (Figure 8D, *p*>0.05).

**Figure 7:**
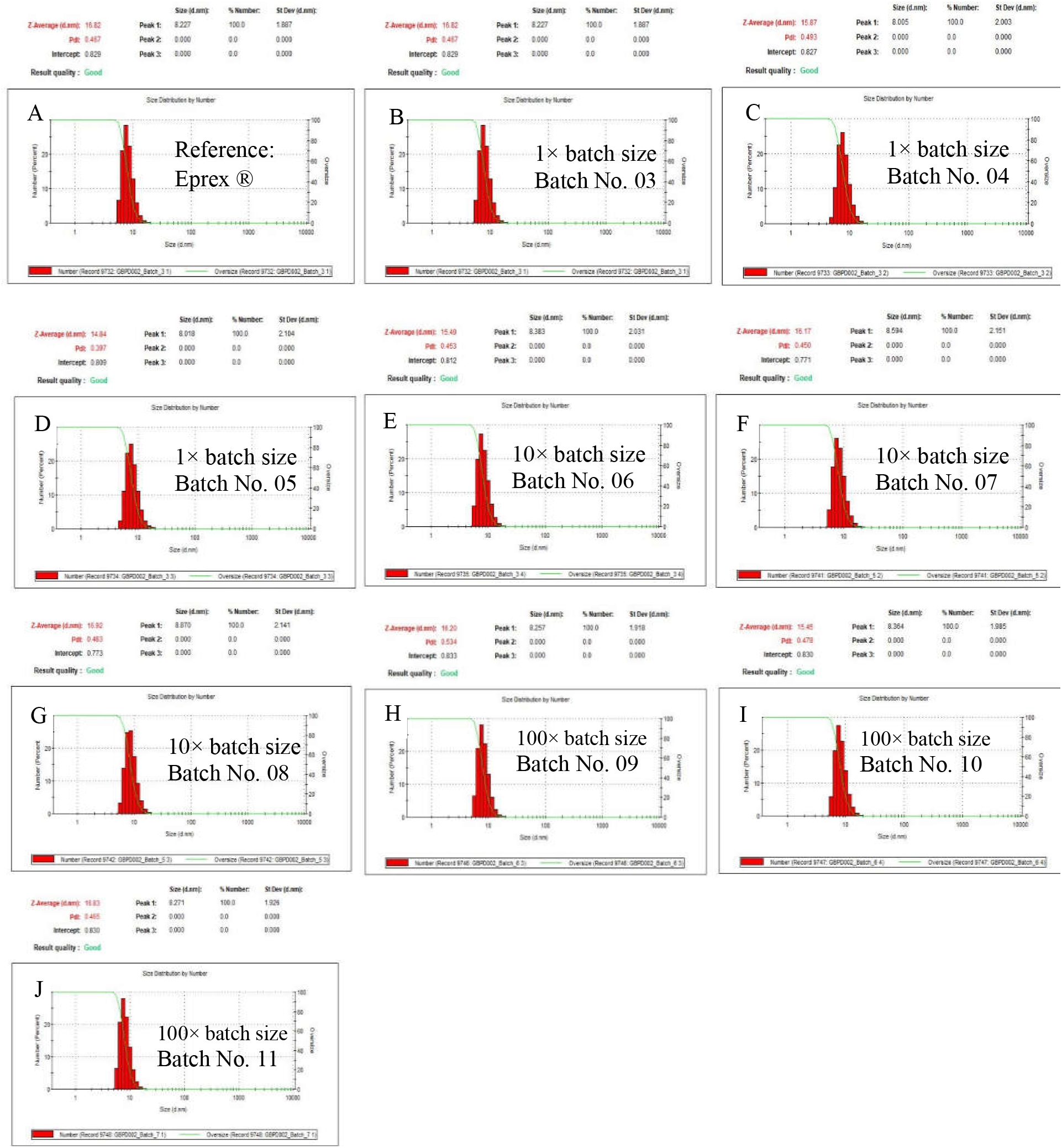
Comparative analysis of particle size distribution between reference product Eprex^®^ and different batches of GBPD002. (**A**), Size distribution of reference product Eprex^®^; **(B, C, D**) size distribution profile of batch no. 03, 04 and 05, respectively (1× batch size); (**E, F** and **G**), size distribution profile of batch no. 06, 07 and 08, respectively (10× batch size); (**H, I** and **J**), size distribution profile of batch no. 06, 07 and 08, respectively (100× batch size).

**Figure 8:**
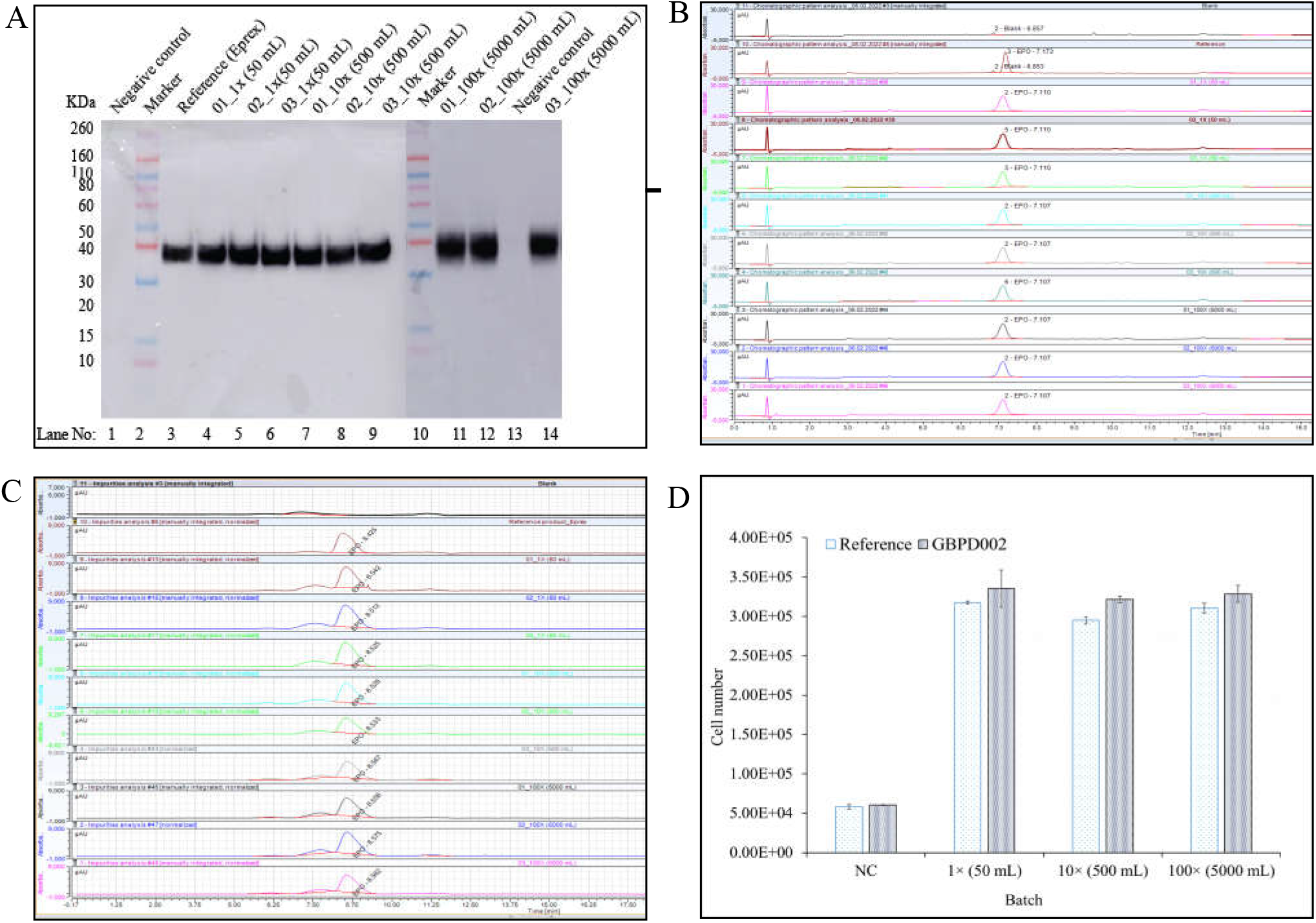
Comparative analysis of QTPP for Eprex^®^ and GBPD002. (**A**) Immunochemical properties analysis by western blot, (**B**) Determination of potency by reverse-phase chromatography, (**C**) Impurities profile analysis by size-exclusion chromatography (**D**) Biofunctionality in cell culture assay.

## 4. Discussion

We have taken a systematic pathway (Scheme-1) to address the challenges associated with the unit processes for the purification of EPO, and transforming the process-train to higher batch sizes with the objectives of meeting regulatory requirements. As per WHO guideline for biologic/biosimilar drug evaluation, the quality attributes of a biologics can be furnished into 3 categories, *viz*., (i) very high critical, (ii) high critical, and (iii) low critical [45]. Amino acid sequence, glycan analysis, biological activity, and immunochemical identity are considered very high critical quality attributes. Higher order structure, isoform distribution, insoluble aggregates, high molecular weight aggregates, protein content, host cell proteins, host cell DNA and receptor binding are considered highly critical quality attributes. Deamination, oxidation, and truncation are considered low critical quality attributes. The amino acid is produced based on the stably integrated relevant DNA of the qualified master cell bank (MCB), and we have found the expressed rhEPO indeed contains all 165 amino acids by LC-MS/MS (data not shown). The MCB has been characterized for 50 consecutive passages over 160 days (data not shown). Therefore, it is highly unlikely that the protein sequence is compromised in current study, where we have used a 2-weeks culture of qualified seeds from the qualified working cell bank (WCB). Instead of amino acid sequencing, therefore, we have used Western blot analysis with a reference product, and similar band pattern and molecular mobility was considered as one of the relevant QTPP. Indeed, Western blot data revealed highly similar single molecular entity like the reference product for all batches from all scales.

Glycosylation profile has been found important for protein folding that affects protein stability and receptor interaction. We did not consider glycoform as a QTPP parameter for this study due to the following reasons. Firstly, though glycoforms analysis are important and may be required for batch release but in practical aspect it is a non-critical parameter for our study as described follows. It has been found that several different EPO-glycoforms were present in circulating systems of human [46], and the sugar profiles of human serum EPO are significantly different from the profiles of rhEPO [47]. Despite having differences in sugar profile, there were no significant differences reported for *in vivo* biofunctionality for several preparations (12 different sources) of rhEPO [48]. Several clinical studies also did not reveal any significant differences between pharmacokinetic and pharmacodynamic properties for different rhEPO [49 – 55]. Secondly, confident glycoform analysis is a very critical process, and multiple parallel methods are preferred for obtaining confident PTM profiles using mass spectrometry analysis [56, 57]. Further, it has been suggested that the separation of recombinant EPO into pure glycoforms is not feasible even with the most advanced methods due to the heterogeneity of the 3 N-glycans [58]. Thirdly, the objective of the study was not the thorough qualification of a candidate EPO in respect of biosimilarity rather to harvest EPO through a well-characterized and scalable downstream process, and use some suitable analytical methods for analyzing the harvest which may conform with the preset specification of EPO.

We have considered aggregation analysis for the QTPP profile of EPO due to the fact that aggregation mainly occurred during the downstream processing particularly, during and on formulation. EPO aggregation has been reported is critically connected for immunochemistry of rhEPO preparation; it was specifically responsible for the development of EPO-antibody mediated deadly disease condition called pure red cell aplasia (PRCA) [59 – 61]. Due to the criticality of the parameter, we have employed 3 orthologous analysis procedures, *viz*., Western blot, dynamic light scattering and SEC-HPLC for detecting any high molecular aggregates in our final products, and all analyses confirmed that our EPO preparations from all 3 scales did not contain any such macromolecular entity.

Quantity and relevant impurities have been considered highly critical quality attributes, and therefore, we considered these parameters as the components of QTPP in our study. We have used orthologous UHPLC protocols in 2 different methods (RPC and SEC) for analyzing these QTPP parameters. All samples from 3 different scale-size (1×, 10× and 100×) batches generated highly similar chromatograms like those of the reference product. The results suggested indifferences between experimental EPO preparations and references in terms of 2 critical QTPP parameters, *viz*., quantity and purity. Since EPO biofunctionality is a concrete proof of functional significance, we therefore, considered the cell-based biofunctional assay as another parameter for QTPP and found similar results for all batches and reference product. The QTPP profile for all validation and scale-up (1×, 10× and 100×) batches satisfied the regulatory acceptance limit of relevant analyses, and thereby proved that the QbD approach used to develop unit processes are effective. Insignificant levels or minor variations in different data point were not affecting the final product quality. In fact, such types of minor variations are not uncommon for US FDA and EMA approved biosimilars [62]. A recent study compared the quality and batch-to-batch variability of marketed rhEPO reference medicines, Eprex®/Erypo® (Janssen-Cilag, High Wycombe, UK) and NeoRecormon® (epoetin beta; Roche Registration Limited, Welwyn Garden City, UK), and two biosimilars, Binocrit® (Sandoz GmbH, Kundl, Austria) and Retacrit® (Hospira UK Limited, Maidenhead, UK), and found batch-batch minor variability for experimental products [63]. Microbial contamination and pyrogen are another 2 QTPP profiles we have included in our study. These contaminants in parenteral drug preparation are life threatening, and mainly enter into the product through the process therefore these parameters are indispensable from QTPP. Relevant results revealed that all EPO preparations were clear from such contaminants and thereby satisfying relevant QTPP.

Several studies have been reported purification of rhEPO; a snap shot of comparative process analysis has been shown in Table 1. The pros and cons of individual processes have been identified and summarized therein. Relevant studies have reported the purification processes of rhEPO with different combinations of multiple unit processes and conditions but none of them has systematically developed their unit processes that may qualify for regulatory framework. Neither the CPPs were identified nor the operating space for any relevant unit processes were characterized in these studies, and thereby severely limiting the scale-up and scale-down opportunities. Several processes are SEC dependent [28 – 34], and SEC is not economic in biopharmaceutical process since it requires huge amount of resin and larger column to handle bulk sample. It also increases process time since it need to run at a slow flowrate because of increasing delta column pressure. Some processes placed SEC unit process after the RPC unit process (where product is in low pH) [28, 29], which is detrimental to the product since EPO degrades fast in low pH. We have avoided expensive and highly critical SEC process and aligned the other processes in the process train to exploit the physicochemical properties of the output of an antecedent unit process as the input for the subsequent unit process, which made the process cost effective and easily scalable. Many processes did not mention virus inactivation and specific step for virus reduction as well as elimination of microbial and pyrogenic contaminants [30 – 34], which is a big concern for regulatory framework to ensure QTPP and patient safety. Our process includes all relevant steps that make the end product safer to patients. Several processes were not aligned with regulatory guidelines since the final product were not in formulated forms and they did not include impurity profile and viral and microbial decontamination processes [35 – 43]. In 2017, Bandi *et al*. has claimed 98% purity for their process but this process has lots of drawbacks. It is associated with at least 3 buffer exchange steps (in between AEC and CEC, CEX and AFC, and AFC and AEX), totaling 10 steps including SEC steps, which is a very expensive and lengthy process though this process is not completed up to formulation step [44].

Our process includes 8 steps to obtain ready-to-fill EPO preparations from the filtered media of upstream satisfying QTPP in line of regulatory framework. Identification of CPPs by DoE in our process made it efficiently convenient to scale-up and validate. After completion of validation, variations between the individual process steps for 1×, 10× and 100× batches were insignificant (p>0.05), as well as the variations within the batches were also insignificant (p>0.05), which have suggested that data were reproduced for 1×, 10× and 100× batches. Collectively, our study provided a scalable and controllable downstream process for purification of drug-quality rhEPO to satisfy regulatory requirements at a competitive cost.

## 5. Conclusion

Here, we presented a systematic approach to develop a well-characterized and scalable purification process of EPO preparation for pharmaceutical use. The operation boundaries for each unit process were validated in 2 steps (10× and 100×) of scale-up process resulting 100× batch size, which provides a tremendous opportunity for seamless integration of the process-train in industrial scale. As a result, EPO can be manufactured in a faster time and cost-effective manner. The process development and validation pathway presented here can be applied to many other such life-saving pharmaceutical products, and thereby will facilitate the relevant downstream process management and promote the availability of cost-effective products to the global community.

## Author contribution

Kakon Nag and Naznin Sultana conceptualized the project. Md. Enamul Haq Sarker, Samir Kumar, Md. Jikrul Islam, Md. Maksusdur Rahman Khan and Md. Mashfiqur Rahman Chowdhury designed experiments and analyzed data. Md. Maksusdur Rahman Khan and Rony Roy contributed to cell line development, and Md. Mashfiqur Rahman Chowdhury contributed to cell bank development and scale-up of the cell culture. Md. Enamul Haq Sarker, Samir Kumar, Sourav Chakraborty and Habiba Khan performed process development and manufacturing steps. Md. Jikrul Islam and Ratan Roy performed quality control experiments. Kakon Nag, Naznin Sultana, Md. Enamul Haq Sarker, Samir Kumar, Sourav Chakraborty and Md. Jikrul Islam wrote the manuscript. Bipul Kumar Biswas, Md. Emrul Hasan Bappi and Mohammad Mohiuddin assured the quality management system of relevant activities.

## Institutional review board statement

Not applicable.

## Informed consent statement

Not applicable.

## Data availability statement

The data that support the findings of this study are available within the article and its Supplementary document file, or are available from the corresponding author upon reasonable request.

## Acknowledgements

The study was funded by Globe Biotech Limited. We thank Md. Harunur Rashid, the chairman of Globe Pharmaceuticals Group of Companies, Ahmed Hossain, Md. Mamunur Rashid, Md. Shahiduddin Alamgir and Abdullah Al Rashid, the directors of Globe Pharmaceuticals Group of Companies for their continuous support and encouragement. We also thank Md. Raihanul Hoque, Dibakor Paul, Biplob Biswas, Alok Sutradhar, Mithun Kumar Nag, Zahir Uddin Babor, G.M. Sajib Hasan and Mijanur Rahman for their support for facility, materials and information management system.

---

**Scheme 1:**
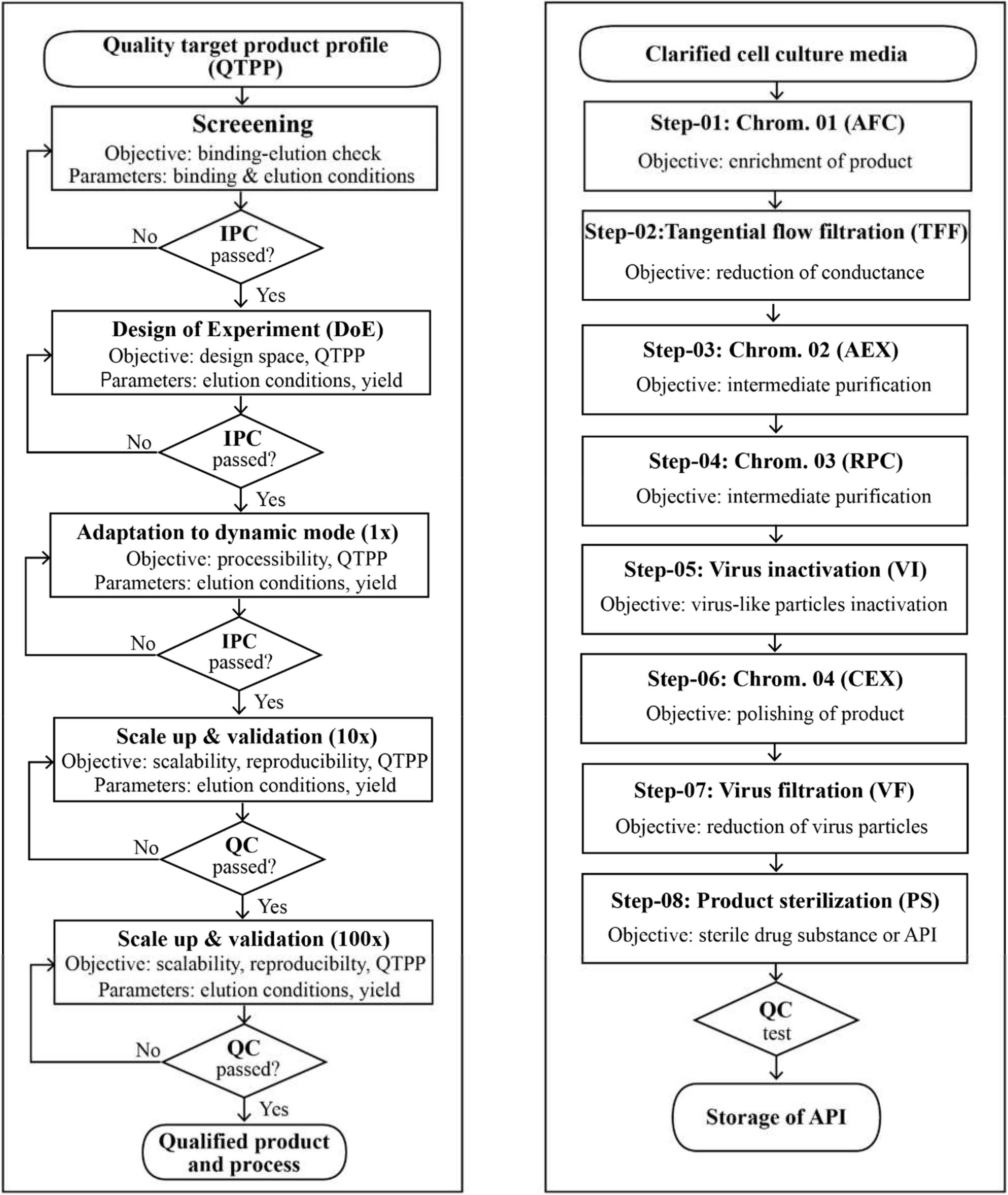
Process development decision tree (**A**) and process flow (**B**) for EPO downstream processing up to ready-to-fill formulation.

