## Supplementary information for "A Cost-effective Purification Process for Erythropoietin Biosimilar through Downstream Process Engineering"

Table 1: DoE design conditions of AFC process step along with recovery %

| Run order | Run No. | pH | [NaCl]mM | %Recovery |
| --- | --- | --- | --- | --- |
| 16 | 1 | 9.4 | 2000 | 41 |
| 13 | 2 | 5.9 | 2000 | 32 |
| 15 | 3 | 7.7 | 2000 | 60 |
| 11 | 4 | 7.6 | 1595 | 95 |
| 3 | 5 | 7 | 1000 | 32 |
| 7 | 6 | 5.4 | 1340 | 36 |
| 12 | 7 | 6.2 | 1645 | 70 |
| 2 | 8 | 7.0 | 1000 | 29 |
| 8 | 9 | 9.3 | 1585 | 55 |
| 4 | 10 | 9.4 | 1174 | 50 |
| 5 | 11 | 9.4 | 1174 | 49 |
| 10 | 12 | 7.6 | 1595 | 91 |
| 1 | 13 | 5.6 | 1000 | 32 |
| 6 | 14 | 8 | 1249 | 56 |
| 14 | 15 | 5.9 | 2000 | 50 |
| 9 | 16 | 7.6 | 1595 | 93 |

Table 2: DoE design conditions of AEX process step along with recovery %

| Run order | Run | pH | [NaCl]mM | %Recovery |
| --- | --- | --- | --- | --- |
| 12 | 1 | 5 | 440 | 24 |
| 7 | 2 | 6.7 | 270 | 40 |
| 4 | 3 | 8.4 | 100 | 10 |
| 13 | 4 | 5 | 440 | 21 |
| 1 | 5 | 5 | 100 | 12 |
| 11 | 6 | 6.4 | 410 | 42 |
| 15 | 7 | 7.5 | 500 | 25 |
| 10 | 8 | 9 | 356 | 30 |
| 16 | 9 | 7.5 | 500 | 23 |
| 14 | 10 | 9 | 500 | 24 |
| 9 | 11 | 5 | 272 | 21 |
| 5 | 12 | 8.2 | 240 | 35 |
| 2 | 13 | 6.7 | 100 | 14 |
| 6 | 14 | 6.7 | 270 | 41 |
| 8 | 15 | 6.7 | 270 | 42 |
| 3 | 16 | 8.4 | 100 | 10 |

Table 3: DoE design conditions of RPC process step along with recovery %

| Run order | Run | pH | %Acetonitrile | %Recovery |
| --- | --- | --- | --- | --- |
| 7 | 1 | 2.5 | 41 | 57 |
| 11 | 2 | 2.7 | 50 | 52 |
| 16 | 3 | 1.8 | 100 | 10 |
| 6 | 4 | 2.2 | 36 | 25 |
| 2 | 5 | 1.5 | 7 | 0 |
| 8 | 6 | 1.5 | 41 | 13 |
| 9 | 7 | 2.7 | 50 | 43 |
| 10 | 8 | 2.7 | 50 | 45 |
| 13 | 9 | 2.7 | 98 | 12 |
| 5 | 10 | 2.8 | 24 | 8 |
| 12 | 11 | 3.5 | 59 | 8 |
| 15 | 12 | 1.8 | 100 | 13 |
| 3 | 13 | 3.5 | 17 | 3 |
| 14 | 14 | 3.5 | 100 | 5 |
| 1 | 15 | 2.2 | 5 | 0 |
| 4 | 16 | 3.5 | 17 | 4 |

Table 4: DoE design conditions of CEX process step along with recovery %

| Run order | Run | pH | Buffer volume (CV) | %Recovery |
| --- | --- | --- | --- | --- |
| 13 | 1 | 2 | 7 | 0 |
| 10 | 2 | 4.2 | 4.4 | 25 |
| 12 | 3 | 5.9 | 4.9 | 60 |
| 8 | 4 | 2 | 4.4 | 0 |
| 7 | 5 | 7 | 3.2 | 52 |
| 11 | 6 | 4.2 | 4.4 | 15 |
| 6 | 7 | 7 | 3.2 | 55 |
| 9 | 8 | 4.2 | 4.4 | 17 |
| 5 | 9 | 3.8 | 2.3 | 12 |
| 1 | 10 | 6 | 4 | 98 |
| 16 | 11 | 6.3 | 7 | 60 |
| 4 | 12 | 2 | 1.8 | 0 |
| 14 | 13 | 4.1 | 7 | 19 |
| 2 | 14 | 5.5 | 4 | 97 |
| 3 | 15 | 2 | 1.8 | 0 |
| 15 | 16 | 6.3 | 7 | 62 |

Table 5: Starting materials quantity, volume and concentration for 50 ml (1×) batch size.

| Batch No. | Initial quantity  (mg) | Sample volume (ml) | Initial concentration (mg/ml) |
| --- | --- | --- | --- |
| 01 | 53 | ~ 50 | 1.06 |
| 02 | 45 | ~ 50 | 0.9 |
| 03 | 55 | ~ 50 | 1.1 |
| 04 | 50 | ~ 50 | 1.0 |
| 05 | 54 | ~ 50 | 1.08 |

Table 6: AFC process adaptation data in dynamic mode for 50 ml (1×) batch size.

| Batch No. | Elution buffer pH | Elution buffer conductance  (mS/cm) | Eluted sample pH | Eluted sample conductance  (mS/cm) | Eluted sample quantity (mg) | Initial quantity (mg) | % Yield |
| --- | --- | --- | --- | --- | --- | --- | --- |
| 01 | 7.45 | 123.14 | 7.43 | 122.60 | 42.96 | 53.00 | 81.06 |
| 02 | 7.24 | 122.21 | 7.21 | 125.25 | 38.56 | 45.00 | 85.69 |
| 03 | 7.40 | 125.32 | 7.39 | 124.80 | 44.13 | 55.00 | 80.24 |
| 04 | 7.56 | 121.61 | 7.52 | 122.64 | 39.68 | 50.00 | 79.36 |
| 05 | 7.52 | 124.82 | 7.22 | 123.61 | 42.78 | 54.00 | 79.22 |
| Average | 7.43 | 123.42 | 7.35 | 123.78 | 41.62 | 51.40 | 81.11 |
| STDEV | ± 0.12 | ±1.61 | ±0.14 | ±1.22 | ±2.38 | ±4.04 | ±2.66 |

Table 7: Buffer exchange process adaptation data in dynamic mode for 50 ml (1×) batch size.

| Batch No. | Retentate sample pH | Retentate sample conductance (mS/cm) | Retentate sample quantity (mg) | Initial quantity (mg) | % Yield |
| --- | --- | --- | --- | --- | --- |
| 01 | 7.41 | 2.16 | 42.51 | 42.96 | 98.95 |
| 02 | 7.22 | 2.59 | 38.23 | 38.56 | 99.14 |
| 03 | 7.40 | 2.32 | 43.41 | 44.13 | 98.37 |
| 04 | 7.54 | 2.83 | 39.12 | 39.68 | 98.59 |
| 05 | 7.19 | 2.76 | 42.18 | 42.78 | 98.60 |
| Average | 7.35 | 2.53 | 41.09 | 41.62 | 98.73 |
| STDEV | ±0.15 | ±0.29 | ±2.27 | ±2.38 | ±0.31 |
| Acceptance limit | 7.35±0.15 | 2.53±0.29 | - | - | 98.73±0.31 |

Table 8: AEX process adaptation data in dynamic mode for 50 ml (1×) batch size.

| Batch No. | Elution buffer pH | Elution buffer conductance  (mS/cm) | Eluted sample pH | Eluted sample conductance  (mS/cm) | Eluted sample quantity (mg) | Initial quantity (mg) | % Yield |
| --- | --- | --- | --- | --- | --- | --- | --- |
| 01 | 7.12 | 123.16 | 7.10 | 24.21 | 17.32 | 42.51 | 40.74 |
| 02 | 7.16 | 121.43 | 6.91 | 25.26 | 15.03 | 38.23 | 39.31 |
| 03 | 6.86 | 122.61 | 7.08 | 24.61 | 18.12 | 43.41 | 41.74 |
| 04 | 6.94 | 125.11 | 6.83 | 24.86 | 16.21 | 39.12 | 41.44 |
| 05 | 7.18 | 123.24 | 6.95 | 26.01 | 17.63 | 42.18 | 41.80 |
| Average | 7.05 | 123.11 | 6.97 | 24.99 | 16.86 | 41.09 | 41.01 |
| STDEV | ±0.14 | ±1.33 | ±0.11 | ±0.69 | ±1.24 | ±2.27 | ±1.03 |

Table 9: RPC process adaptation data in dynamic mode for 50 ml (1×) batch size.

| Batch No. | Elution buffer pH | Acetonitrile % in elution buffer | Eluted sample pH | Acetonitrile % in sample | Eluted sample quantity (mg) | Initial quantity (mg) | % Yield |
| --- | --- | --- | --- | --- | --- | --- | --- |
| 01 | 2.34 | 95 | 2.20 | 50 | 8.51 | 17.32 | 49.13 |
| 02 | 2.54 | 95 | 2.00 | 48 | 7.82 | 15.03 | 52.03 |
| 03 | 2.41 | 95 | 2.29 | 52 | 9.16 | 18.12 | 50.55 |
| 04 | 2.36 | 95 | 2.16 | 51 | 8.69 | 16.21 | 53.61 |
| 05 | 2.5 | 95 | 2.21 | 53 | 9.21 | 17.63 | 52.24 |
| Average | 2.43 | 95 | 2.17 | 50.80 | 8.68 | 16.86 | 51.51 |
| STDEV | ±0.09 | 0 | ±0.11 | ±1.92 | ±0.57 | ±1.24 | ±1.72 |

Table 10: CEX process adaptation data in dynamic mode for 50 ml (1×) batch size.

| Batch No. | Elution buffer pH | Elution buffer volume (CV) | Eluted sample pH | Eluted sample quantity (mg) | Initial quantity (mg) | % Yield |
| --- | --- | --- | --- | --- | --- | --- |
| 01 | 7.22 | 3.5 | 5.71 | 7.65 | 8.51 | 89.86 |
| 02 | 6.8 | 3.5 | 5.82 | 6.25 | 7.82 | 79.92 |
| 03 | 7.15 | 3.5 | 5.68 | 8.47 | 9.16 | 92.45 |
| 04 | 6.91 | 3.5 | 5.89 | 6.60 | 8.69 | 75.95 |
| 05 | 7.26 | 3.5 | 5.93 | 8.39 | 9.21 | 91.11 |
| Average | 7.07 | 3.5 | 5.81 | 7.47 | 8.68 | 85.86 |
| STDEV | ±0.20 | 0 | ±0.11 | ±1.02 | ±0.57 | ±7.42 |

Table 11: Virus filtration process data for 50 ml (1×) batch size.

| Batch No. | Filtered  sample pH | Filtered  sample conductance  (mS/cm) | Filtered sample quantity (mg) | Initial quantity (mg) | % Yield |
| --- | --- | --- | --- | --- | --- |
| 01 | 7.20 | 12.5 | 7.56 | 7.65 | 98.86 |
| 02 | 7.18 | 12 | 6.18 | 6.25 | 98.91 |
| 03 | 7.09 | 13.19 | 8.31 | 8.47 | 98.14 |
| 04 | 7.22 | 12.63 | 6.58 | 6.60 | 99.80 |
| 05 | 6.92 | 12.39 | 8.25 | 8.39 | 98.29 |
| Average | 7.12 | 12.54 | 7.38 | 7.47 | 98.80 |
| STDEV | ±0.12 | ±0.43 | ±0.97 | ±1.02 | ±0.65 |
| Acceptance limit | - | - | - | - | 98.80±0.65 |

Table 12: Sterile filtration process data for 50 ml (1×) batch size.

| Batch No. | Filtered  sample pH | Filtered  sample conductance  (mS/cm) | Filtered sample quantity (mg) | Initial quantity (mg) | % Yield |
| --- | --- | --- | --- | --- | --- |
| 01 | 7.22 | 12.45 | 7.42 | 7.56 | 98.15 |
| 02 | 7.18 | 12.11 | 6.16 | 6.18 | 99.69 |
| 03 | 7.1 | 13.16 | 8.25 | 8.31 | 99.27 |
| 04 | 7.21 | 12.59 | 6.50 | 6.58 | 98.84 |
| 05 | 6.93 | 12.41 | 8.10 | 8.25 | 98.21 |
| Average | 7.13 | 12.54 | 7.29 | 7.38 | 98.83 |
| STDEV | ±0.12 | ±0.39 | ±0.93 | ±0.97 | ±0.67 |
| Acceptance limit | - | - | - | - | 98.83±0.67 |

Table 13: Overall yield data of 50 ml (1×) batch size.

| Batch No. | Initial quantity  (mg) | Final quantity  (mg) | % Yield |
| --- | --- | --- | --- |
| 01 | 53.00 | 7.42 | 14.00 |
| 02 | 45.00 | 6.16 | 13.69 |
| 03 | 55.00 | 8.25 | 15.00 |
| 04 | 50.00 | 6.50 | 13.00 |
| 05 | 54.00 | 8.10 | 15.00 |
| Average | - | - | 14.14 |
| STDEV | - | - | ±0.87 |
| Acceptance limit | - | - | 14.14±0.87 |

Table 14: Validation of 500 ml (10×) of batch size.

| Parameter | Batch  06 | % Yield | Batch 07 | % Yield | Batch 08 | % Yield | Acceptance limit of % yield | Observation |
| --- | --- | --- | --- | --- | --- | --- | --- | --- |
| Starting sample volume (ml) | 500 | - | 500 | - | 500 | - | - | - |
| Starting sample concentration (mg/ml) | 1.04 | - | 1.01 | - | 0.98 | - | - | - |
| Starting sample quantity (mg) | 520.00 | - | 505.00 | - | 489.00 | - | - | - |
| Quantity of AFC elute (mg) | 417.87 | 80.36 | 414.76 | 82.13 | 405.67 | 82.82 | ≥ 80% | Within limit |
| Quantity of TFF retentate (mg) | 411.73 | 98.53 | 408.99 | 98.61 | 400.93 | 99.37 | 98.73±0.31% | Within limit |
| Quantity of AEX elute (mg) | 164.40 | 39.93 | 167.44 | 40.94 | 169.87 | 39.43 | ≥ 40% | Within limit |
| Quantity of RPC elute (mg) | 82.55 | 50.21 | 85.11 | 50.83 | 85.23 | 49.95 | ≥ 40% | Within limit |
| Quantity of CEX elute (mg) | 75.01 | 90.87 | 73.36 | 85.19 | 69.36 | 82.20 | ≥ 80% | Within limit |
| Quantity of VF sample (mg) | 73.74 | 98.31 | 72.01 | 98.16 | 68.71 | 98.88 | 98.80±0.65% | Within limit |
| Quantity of SF sample (mg) | 72.62 | 98.48 | 71.06 | 98.68 | 67.98 | 99.41 | 98.83±0.67% | Within limit |
| Overall % yield |  | 13.97 |  | 14.07 |  | 13.90 | 14.14±0.87% | Within limit |

Table 15: Validation of 5000 ml (100×) of batch size.

| Parameter | Batch 09 | % Yield | Batch  10 | % Yield | Batch  11 | % Yield | Acceptance limit of % yield | Decision |
| --- | --- | --- | --- | --- | --- | --- | --- | --- |
| Starting sample volume (ml) | 5000 | - | 5000 | - | 5000 | - | - | - |
| Starting sample concentration (mg/ml) | 1.09 | - | 0.99 | - | 1.08 | - | - | - |
| Starting sample quantity (mg) | 5445 | - | 4950 | - | 5376 | - | - | - |
| Quantity of AFC elute (mg) | 4450.02 | 81.73 | 4579.17 | 92.51 | 4493.78 | 83.59 | ≥ 80% | Within limit |
| Quantity of TFF retentate (mg) | 4406.43 | 99.02 | 4550.16 | 99.37 | 4480.27 | 99.70 | 98.73±0.31% | Within limit |
| Quantity of AEX elute (mg) | 1812.33 | 41.13 | 1794.21 | 39.43 | 1837.75 | 41.02 | ≥ 40% | Within limit |
| Quantity of RPC elute (mg) | 910.318 | 50.23 | 896.2 | 49.95 | 937.28 | 51.00 | ≥ 40% | Within limit |
| Quantity of CEX elute (mg) | 747.91 | 82.16 | 736.67 | 82.20 | 735.73 | 78.50 | ≥ 80% | Within limit |
| Quantity of VF sample (mg) | 733.71 | 98.10 | 728.39 | 98.88 | 720.87 | 97.98 | 98.80±0.65% | Within limit |
| Quantity of SF sample (mg) | 724.06 | 98.68 | 724.11 | 99.41 | 715.34 | 99.23 | 98.83±0.67% | Within limit |
| Overall % yield | - | 13.30 | - | 14.62 | - | 13.31 | 14.14±0.87% | Within limit |

Table 16: Analytical findings of three different validation batches (1×, 10× and 100×).

| **Test parameters** | **Batch_1× (50 mL)** | | | **Batch_10× (500 mL)** | | | **Batch_100× (5000 mL)** | | |
| --- | --- | --- | --- | --- | --- | --- | --- | --- | --- |
|  | 01 | 02 | 03 | 01 | 02 | 03 | 01 | 02 | 03 |
| ***General test*** |  |  |  |  |  |  |  |  |  |
| Appearance | Pass | Pass | Pass | Pass | Pass | Pass | Pass | Pass | Pass |
| pH (7.0 ± 0.30) | 6.96 | 6.86 | 6.99 | 6.98 | 6.97 | 6.99 | 6.88 | 6.97 | 6.85 |
| Extractable volume | Pass | Pass | Pass | Pass | Pass | Pass | Pass | Pass | Pass |
| Uniformity of dosage units (L1≤15) | 6.7 | 6.3 | 7.7 | 6.3 | 6.4 | 6.3 | 6.3 | 6.4 | 6.7 |
| Sub-visible particles |  |  |  |  |  |  |  |  |  |
| >10 µm | Pass | Pass | Pass | Pass | Pass | Pass | Pass | Pass | Pass |
| >25 µm | Pass | Pass | Pass | Pass | Pass | Pass | Pass | Pass | Pass |
| ***Identification*** | | | | | | | | | |
| Isoform pattern | Pass | Pass | Pass | Pass | Pass | Pass | Pass | Pass | Pass |
| Molecular mass (KDa) | ~40 | ~40 | ~40 | ~40 | ~40 | ~40 | ~40 | ~40 | ~40 |
| Peptide mapping | Pass | Pass | Pass | Pass | Pass | Pass | Pass | Pass | Pass |
| Glycosylation pattern | Pass | Pass | Pass | Pass | Pass | Pass | Pass | Pass | Pass |
| ***Impurity*** | | | | | | | | | |
| Aggregation | Pass | Pass | Pass | Pass | Pass | Pass | Pass | Pass | Pass |
| Host cell DNA (ng) | <10 | <10 | <10 | <10 | <10 | <10 | <10 | <10 | <10 |
| Host cell protein (ng) | *bdl | *bdl | *bdl | *bdl | *bdl | *bdl | *bdl | *bdl | *bdl |
| ***Assay*** |  |  |  |  |  |  |  |  |  |
| *In-vitro* assay | Pass | Pass | Pass | Pass | Pass | Pass | Pass | Pass | Pass |
| *In-vivo* assay  (80-125%) | 104 | 104 | 105 | 106 | 110 | 108 | 106 | 104 | 107 |
| Receptor binding | Pass | Pass | Pass | Pass | Pass | Pass | Pass | Pass | Pass |
| Potency (80-125%) | 104 | 107 | 105 | 106 | 105 | 107 | 104 | 103 | 105 |
| Sterility | Pass | Pass | Pass | Pass | Pass | Pass | Pass | Pass | Pass |
| Bacterial endotoxins (EU/mL) | 0.4 | 0.3 | 0.4 | 0.2 | 0.3 | 0.4 | 0.5 | 0.3 | 0.2 |

*bdl=Below detection limit

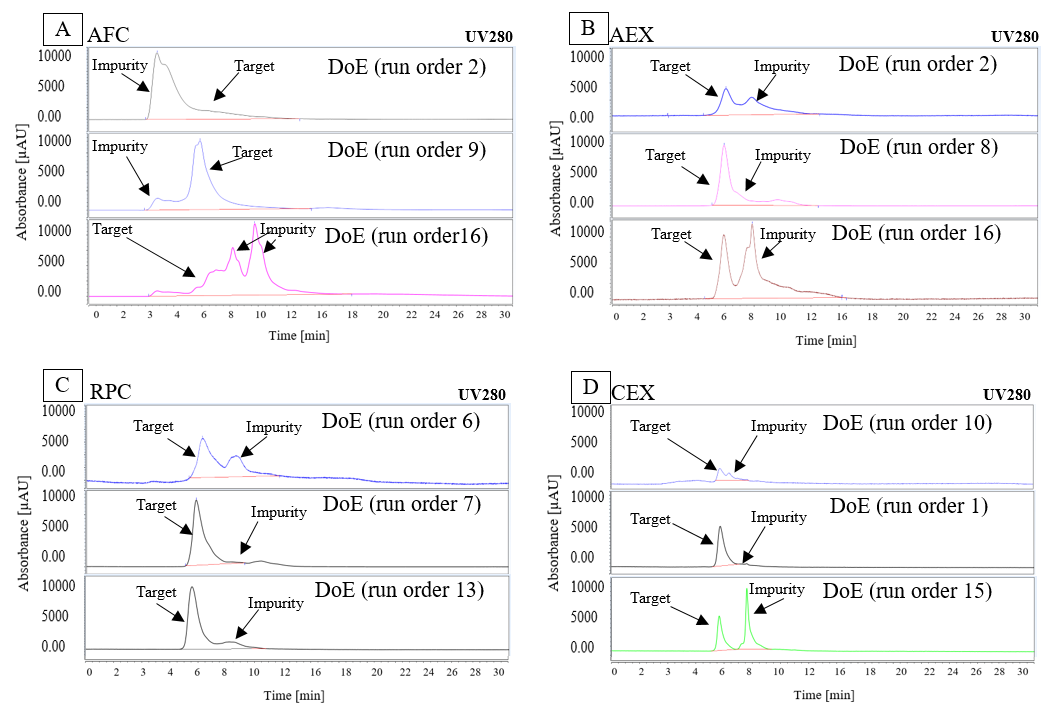
Figure

**Figure 1**: Size exclusion chromatography (SEC) HPLC analysis of representative samples for different process steps originated from DoE recommended runs. (A) SEC chromatograms shows target product and impurities profile of AFC samples for run order 2,9 and 16 and run order 9 shows higher recovery with least impurities than run order 2 and 16, (B) SEC chromatograms shows target product and impurities profile of AEX samples for run order 8,2 and 16 and run order 8 shows higher recovery with least impurities than run order 2 and 16, (C) SEC chromatograms shows target product and impurities profile of RPC samples for run order 6,7 and 13 and run order 7 shows higher recovery with least impurities than run order 7 and 13, (D) SEC chromatograms shows target product and impurities profile of CEX samples for run order 10,1 and 15 and run order 1 shows higher recovery with least impurities than run order 10 and 15.
